# Long-term stability of neuronal ensembles in mouse visual cortex

**DOI:** 10.1101/2020.10.28.359117

**Authors:** Jesús Pérez-Ortega, Tzitzitlini Alejandre-García, Rafael Yuste

## Abstract

Coactive neuronal ensembles are found in spontaneous and evoked cortical activity and are thought to participate in the internal representation of memories, perceptions, and mental states. In mouse visual cortex, ensembles can be optogenetically imprinted and are causally related to visual percepts, but it is still unknown how stable they are over time. Using two-photon volumetric microscopy, we performed calcium imaging over several weeks of the same neuronal populations in layer 2/3 of visual cortex of awake mice, tracking over time the activity of the same neurons in response to visual stimuli and under spontaneous activity. Only a small number of neurons remained active across days. Analyzing them, we found both stable ensembles, lasting up to 46 days, and transient ones, observed during only one imaging session. The majority of ensembles in visually-evoked activity were stable, whereas in spontaneous activity similar numbers of stable and transient ensembles were found. Among stable ensembles, more than 60 % of neurons still belonged to the same ensemble even after several weeks. These core ensemble cells had stronger functional connectivity than neurons that stopped belonging to the ensemble. Our results demonstrate that spontaneous and evoked neuronal ensembles can last weeks, providing a neuronal mechanism for the long-lasting representation of perceptual states or memories.

## Introduction

Neuronal ensembles, defined as a group of neurons which tend to fire together, are thought to underlie the neural representations of memories, perceptions, thoughts, motor programs, computations, or mental states (Lorente de No, 1938; Hebb, 1949; Cossart et al., 2003; Ikegaya et al., 2004; Sasaki et al., 2007, Buzsáki, 2010; Shepherd & Grillner, 2010; Yuste, 2015; Stringer et al. 2019a; Carrillo-Reid & Yuste, 2020). Using two-photon calcium imaging, ensembles have been found in visual cortex during spontaneous activity and after visual stimulation (Cossart et al., 2003; Miller et al., 2014; Carrillo-Reid et al., 2015a; Stringer et al. 2019b). Ensembles can also be optogenetically imprinted (Carrillo-Reid et al., 2016) and their activation can lead to behavioral effects, consistent with the hypothesis that they represent perceptual states (Carrillo-Reid et al., 2019; Marshel et al., 2019). If this is the case, one would expect those representations to be preserved over time, to retain learned memories or perception. However, it still unknown how stable neuronal ensembles are. To investigate this, we performed longitudinal calcium imaging experiments using two-photon volumetric microscopy and tracked the responses of same neurons for up to 46 days in the visual cortex of awake mice. Functional connectivity based on neuronal coactivity was used to detect neuronal ensembles and we examined if ensembles were preserved in the following days. During spontaneous and visually-evoked activity we found stable ensembles and transient ensembles, present in only one session. In spontaneous activity, approximately ∼50% of ensembles were stable whereas during evoked activity ∼70% were stable. We also analyzed neuronal participation within stable ensembles over time, finding stable neurons that belong to the same ensemble through days; neurons that stopped responding or became part of other ensembles; and new neurons that enrolled into an existing ensemble. Functional connectivity analysis revealed that stable neurons were more connected than neurons which were eventually lost. Our results reveal long-term stability over several weeks of ensembles constituted mainly by neurons that are more functionally connected.

## Results

### Experimental and analysis rationale

To examine ensembles stability under visually-evoked and spontaneous activity we performed two-photon calcium imaging in layer 2/3 of the visual cortex from transgenic mice (GCaMP6s, n = 4; and GCaMP6f, n = 2) through a cranial window. We head-fixed a mouse in front of a blue screen monitor (Figure 1A) and first recorded the spontaneous neuronal activity using a plain static blue screen (Figure 1B). Then we recorded visually evoked activity by showing 50 times a single-orientation blue drifting gratings stimulus (2 s each) with a plain static blue screen between presentations (at 1 to 5 s random intervals; Figure 1C). For either spontaneous or evoked activity, we recorded 3 sessions each day. In order to track the same neurons across days, volumetric calcium imaging was performed. A reference plane (0 µm) from day 1 was first located, then 2 extra planes 5 µm apart were recorded, above (−5 µm) and below (+5 µm) the reference plane (Figure 1D left). At the end of the imaging, maximum intensity projection frames were created from the 3 planes to assemble a single video per session (Figure 1D right). In these datasets, we identified the regions of interest (ROIs) of neuronal activity and kept neurons with peak signal to noise ratio (PSNR) > 18 dB (Figure 1E left). Calcium signals from each ROI were then deconvolved for spike inference and then thresholded to generate a binarized signal, which we used to build spike rasters (Figure 1E right). We then analyzed all ROIs to build a binary raster (Figure 1F) and recorded the activity of the same neurons on day 2, 10 and day 43 or 46 (Figure 1G — Table 1). Some animals were imaged in day 43 and others in day 46, but for the statistics, we combined the data as a single day 43-46.

**Figure 1.**
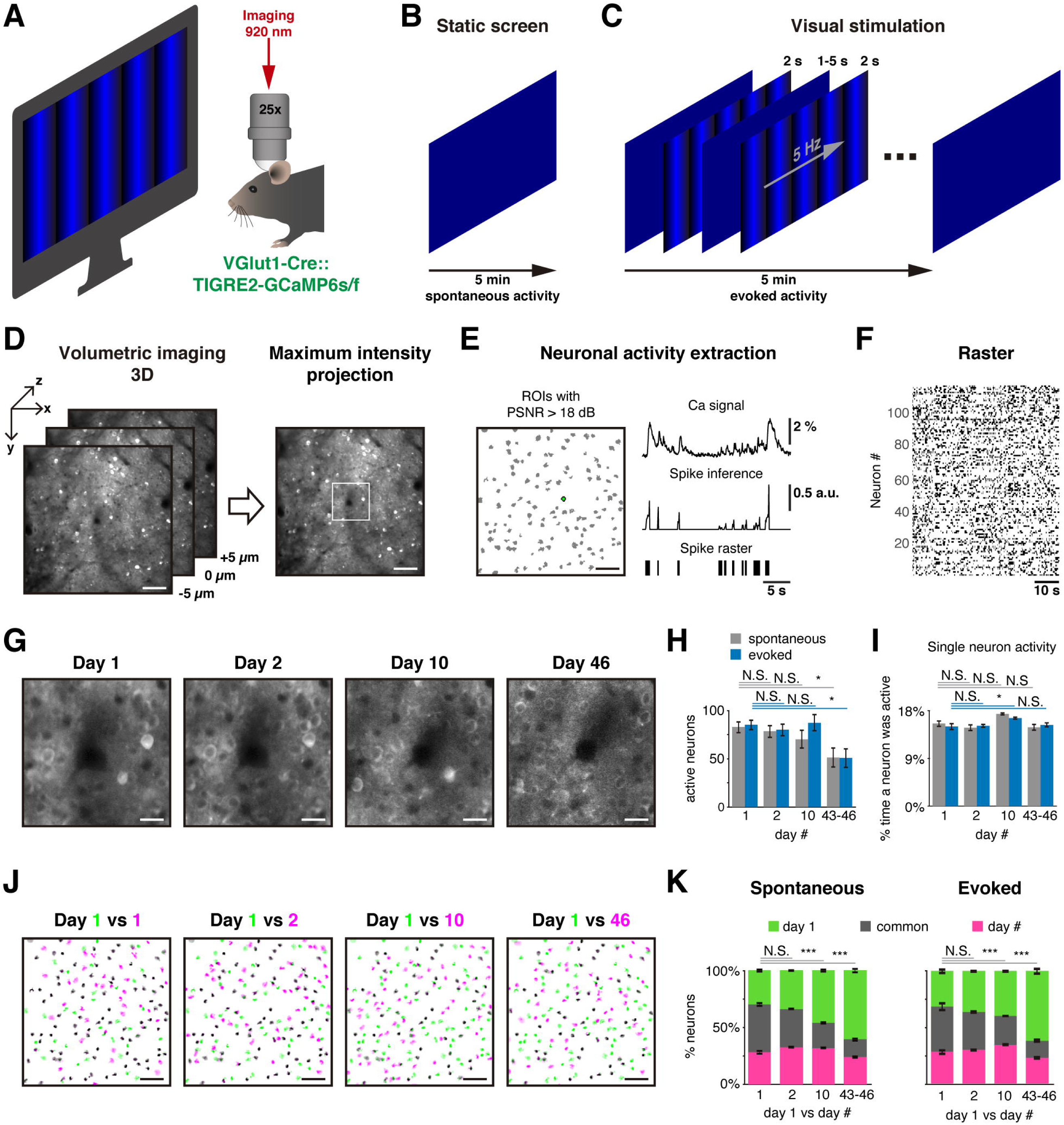
Experimental protocol for longitudinal tracking. A. Experimental set up: a mouse with cranial window is placed on a treadmill in front of a blue screen monitor, and it is head-fixed under the two-photon microscope for calcium imaging recording. B. Static blue screen was used to record spontaneous activity during 5 min, 3 sessions per day, 5 min apart between them. C. Visual stimulation protocol constituted of 50 times of a 2 s single-orientation drifting gratings with a mean static screen between each of them during 1-5 s randomly to record evoked activity for 5 min, 3 sessions per day, 5 min apart between them. D. Strategy to image the same neurons in the field of view in different days: left, a single plane was carefully located in the reference recorded place (day 1), then two extra planes also were recorded separated 5 µm up and down. Three planes were recorded in a period of 81 ms. Right, maximum intensity projection was obtained from the three planes recorded to have a single video per session. Scale bar: 50 µm. E. Left, detection of ROIs (gray shapes) based on Suite2P algorithm, green ROI is used as representative example, scale bar: 50 µm; up right, extraction of Ca signal with PSNR > 18 dB; middle right, spike inference using foopsi algorithm; bottom right, binary signal obtained by thresholding spike inference, which is used to represent the active frames of the neuron. F. Raster plot built with binary signals from active neurons recorded simultaneously. Each row represents the activity of a single neuron, black dots represent the activity of the neuron. G. Center of an example image at 4X zoom (white square on D right) recorded in the same location up to 46 days after the first day of recording. Note image at day 46 is noisier than first days. Scale bar: 12.5 µm. H. Count of active neurons in different days. The number of active neurons identified on day 1 decreased significantly on day 43-46 during spontaneous and evoked activity (p = 0.023 and p = 0.013, respectively). I. Percentage of single neuron activity, i.e. percentage of frames a neuron was active. The average activity of all neurons was around ∼15% on all days during spontaneous and evoked activity. J. Merge of active neuron ROIs from two sessions: first session from day 1 (green in four panels) versus a second single session from same day 1 (5 min later), one session from day 2, 10 and 46 (from left to right respectively, magenta). The intersection of active neurons in both sessions are in gray color. Scale bar: 50 µm. K. Percentage of active neurons between two sessions: first session from day 1 (green) vs second sessions from day 1, 2, 10 and 46 (magenta). The common active neurons (gray) in both sessions were 42 ± 2 % during spontaneous and 37 ± 4 % during evoked activity. There were no significant differences on common active neurons between day 1 and 2, but a significant decrease on days 10 and 43-46 during spontaneous (p = 2×10^−6^ and p = 4×10^−9^, respectively) and evoked activity (p = 2×10^−4^ and p = 4×10^−9^, respectively). Data are presented as mean ± SEM. Kruskal-Wallis test with post hoc Tukey-Kramer: * p < 0.05, ** p < 0.01 and *** p < 0.001. See Figure 1 — Table 2.

We first examined if recording time determined the number of active cells found. As neurons become active at different times, one would expect to capture more active neurons the longer the recording session. At the same time, a very long imaging session would not be practical and could make difficult the focus of our study, which is to identify the change over time of neuronal ensembles. To define an imaging interval that significantly captured the neuronal activity present in the imaged field, pilot experiments and tabulated data across imaging sessions of increasing duration were carried out (Figure 1 — Figure supplement 1). The accumulated number of active neurons reached a plateau after a few minutes. Based on this curve, we reasoned that, with our imaging and analysis pipeline, intervals of 5 minutes would capture the majority of active neurons in the imaged territories, and carried out the rest of the study by imaging spontaneous and evoked activity in 5 minute intervals.

### A minority of neurons remain active across days

We then inquired if the number of active neurons was constant across time, and counted active neurons on different days. On day 1, we found an average of 83 ± 6 and 85 ± 5 (mean ± SEM) active neurons during spontaneous and evoked activity imaging periods, respectively. While the number of active neurons was similar in days 1-10, a significant decrease in the number of active neurons occurred in day 43-46, in both spontaneous and evoked activity (Figure 1H; 51 ± 10 neurons; p = 0.023 and p = 0.013, respectively). This decrease could be due to several factors, including diminishing quality of transgene expression and repeated experimental procedures on the same cortical territory, for example animal manipulation, surgical attachments, microtraumas and laser exposures. On the other hand, the percentage of time that a neuron was active (i.e. the number of frames with activity of a given neuron) did not change, and was 15.5 ± 0.5 % and 15 ± 0.6 % (mean ± SEM), for spontaneous and evoked activity respectively on day 1, with no significance difference (one exception) on the following days (Figure 1I). This indicated that, on an individual neuron basis, the level of neuronal activity remained similar across days. Thus, if a neuron is active on any given day, this activity is similar.

We then explored if active neurons were repeatedly active across days, and analyzed if the neurons that were active on the first session of the first day were also active in later sessions, up to 46 days later (Figure 1J). A minority of neurons were active across days, and their proportion became reduced over time (Figure 1K). Even within day 1, only 42 ± 2 % and 37 ± 4 % (mean ± SEM) of neurons were active in subsequent imaging sessions, 5 minutes later, during spontaneous and evoked activity (gray on Figure 1K). This low number could partly be explained by the fact that not all active neurons were captured in a 5 minute interval (Figure 1 — Figure supplement 1) but also likely are due to the possibility that a significant number of neurons were only active during part of the time. Consistent with this, across days, the percentage of neurons that remained active continued to decrease, with significant decrements on day 10 and 43-46, in both spontaneous and evoked activity (gray on Figure 1K). In spite of this generalized decrease in the number of neurons that remained active, during spontaneous and evoked activity, we found 17 ± 7 and 22 ± 8 (mean ± SEM) neurons on day 1 that were still active 46 days later. We concluded that the neurons that are activated by visual stimuli or spontaneously changed over time, with only a small minority of cells maintaining their responses across weeks.

### Neuronal ensembles identification based on functional connectivity

We then explored if neuronal ensembles were present in the dataset. Given that the majority of the neurons did not remain active across different days, to perform the neuronal ensemble analysis we only analyzed neurons that remained active across sessions. To do so, we built binary raster plots of neurons that remained active across sessions (Figure 2A) and detected ensembles using their functional connectivity (Pérez-Ortega et al., 2016). Specifically, to identify a significant coactivation between every pair of neurons we first generated 1,000 spike raster surrogates by a random circular shift in time of the active frames (Figure 2B left). Then we tabulated how many times a given pair of neurons were coactive by chance, and used a 95 % threshold on the cumulative probability from surrogate coactivations (Figure 2B right). A functional connection was assigned if the coactivations were above chance level (p < 0.05). This process was done for every pair of neurons to build a functional network graph (Figure 2C). We then filtered the raster plot removing the activity without significant coactivation and kept only frames with 3 or more active neurons (Figure 2D). Then, treating each frame as a vector (1 frame bin = 81 ms), we computed the Jaccard similarity between every pair of vectors (Figure 2D bottom). In this case, Jaccard similarity indicates the fraction of active neurons common between 2 vectors, i.e. a value of zero means that neurons from one vector are all different from neurons from the other vector, while a value of one means that all neurons between both vectors are the same. To detect ensembles, we searched for repeated patterns of coactivation (or clusters of similar vectors) by performing a hierarchical clustering of all vectors by single linkage, keeping only the most similar vectors (> 2/3 Jaccard similarity, red dotted line on Figure 2E). Similar vectors were clustered by Ward linkage using contrast index to determine the number of groups, i.e. neuronal ensembles (Figure 2F). Finally, we extracted the neuron identity from the ensembles and built ensemble raster plots and spatial maps of them (Figure 2G; see Methods for details).

**Figure 2.**
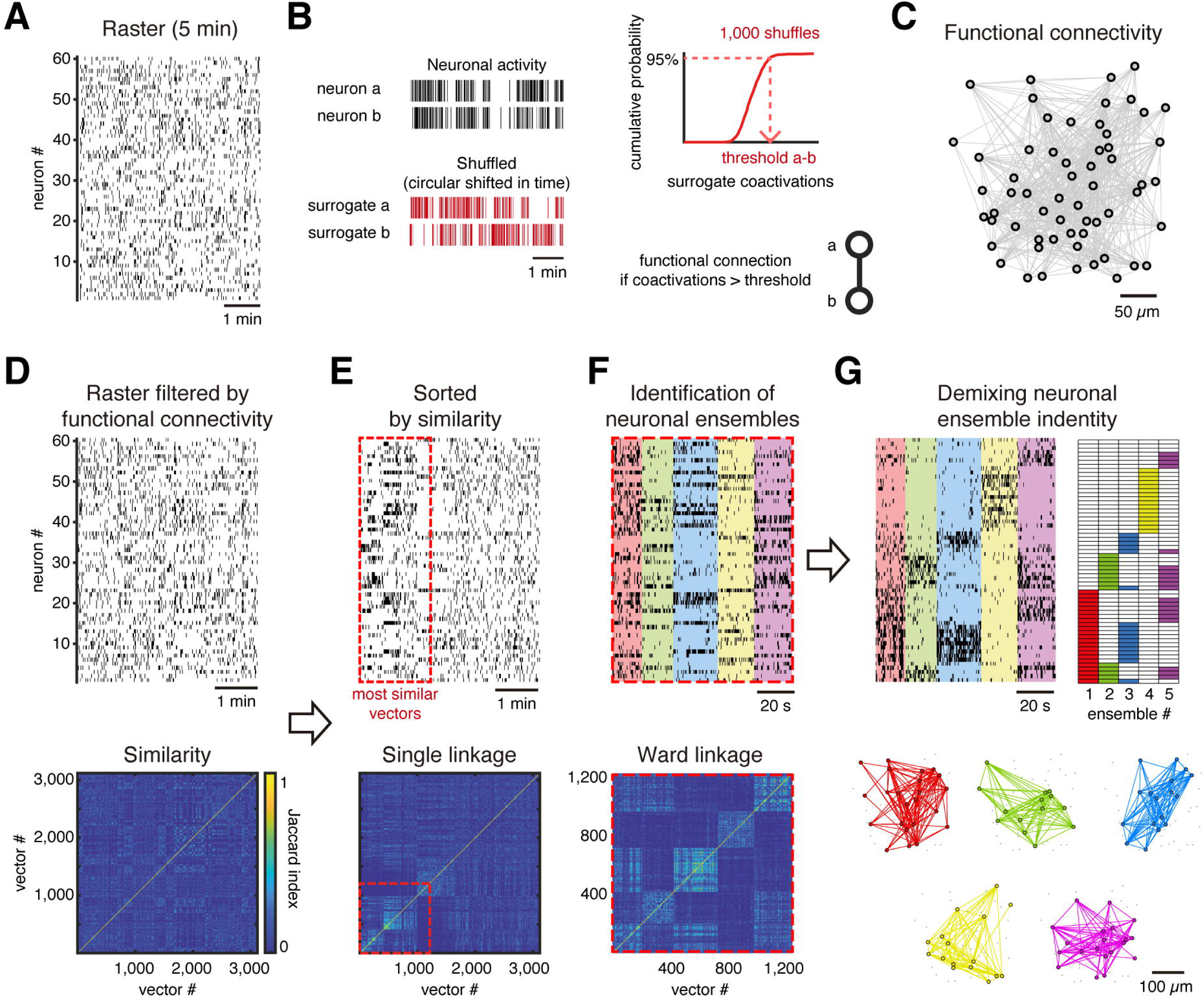
Ensemble identification. C. Example of a raster plot from a single session of spontaneous activity. D. Left, from each pair of neurons *a* and *b* were generated 1,000 surrogates by circular shift of the activity, in a random amount of time, to disrupt the temporal dependency. Right, A cumulative distribution probability of the surrogate coactivations for each pair of neurons is built, which is used to define a threshold of the number of coactivations at 95 % of chance. Then we put a functional connection between those neurons if they reach a significant number of coactivations (p < 0.05), i.e. a number of coactivations bigger than the threshold. E. Functional neuronal network obtained from raster on A, where every node represents a neuron and every link represents a functional connection between neurons. The network is plotted preserving the spatial location of the neurons. F. Top, rebuilt raster from A based on functional connectivity, removing activity from neurons that were active with neurons with no significant coactivity and keeping only frames with 3 or more coactive neurons. Bottom, Jaccard similarity between every column vector (single frame). G. Top, column vectors from raster on D sorted by hierarchical clustering using single linkage based on its Jaccard similarity (bottom). Red dotted line indicates the most similar vectors depicted by thresholding the hierarchical clustering with Jaccard similarity > 2/3. H. Top, most similar vectors on E sorted by hierarchical clustering using Ward linkage based on Jaccard similarity (bottom) and grouped in different ensembles (different color each) based on the contrast index. I. Top left, same sorted vectors on F but here the neurons were sorted depending on their belonging ensemble (top right). Bottom, functional neuronal networks representing the ensembles plotted preserving the spatial location of the neurons.

### Spontaneous and evoked stable neuronal ensembles can last up to 46 days

Using this approach, we identified an average of 4.57 ± 0.14 and 4.63 ± 0.14 (mean ± SEM) ensembles on spontaneous and evoked activity, with no significant differences between them (Figure 2 — Figure supplement 1). We then inquired if ensembles were preserved across days. To do so, we defined as the same neuronal ensemble across days if at least 50 % of its neurons were preserved. This 50% criteria was chosen solely as an operational definition of stability. In comparisons made between 2 imaging sessions from either spontaneous or evoked activity, we found that some ensembles were preserved but others were not (Figure 3A-B). We termed “stable” the ensembles found on day 1 which were preserved in subsequent days, and “transient” all other ensembles. From here on, we focused our analysis on stable ensembles (Figure 3 — Videos 1-2). On day 1, stable ensembles constituted 54 ± 3 % (mean ± SEM) of all ensembles during spontaneous activity, but interestingly were 72 ± 4 % (mean ± SEM) of ensembles in evoked activity (Figure 3C). Similar trends were observed on days 2 and 10. On day 43-46, stable ensembles were around ∼50 % of all ensembles during spontaneous or evoked activity (Figure 3C). This suggested that evoked ensembles were more stable across days 1-10, but are similarly stable as spontaneous ones after 43 days.

**Figure 3.**
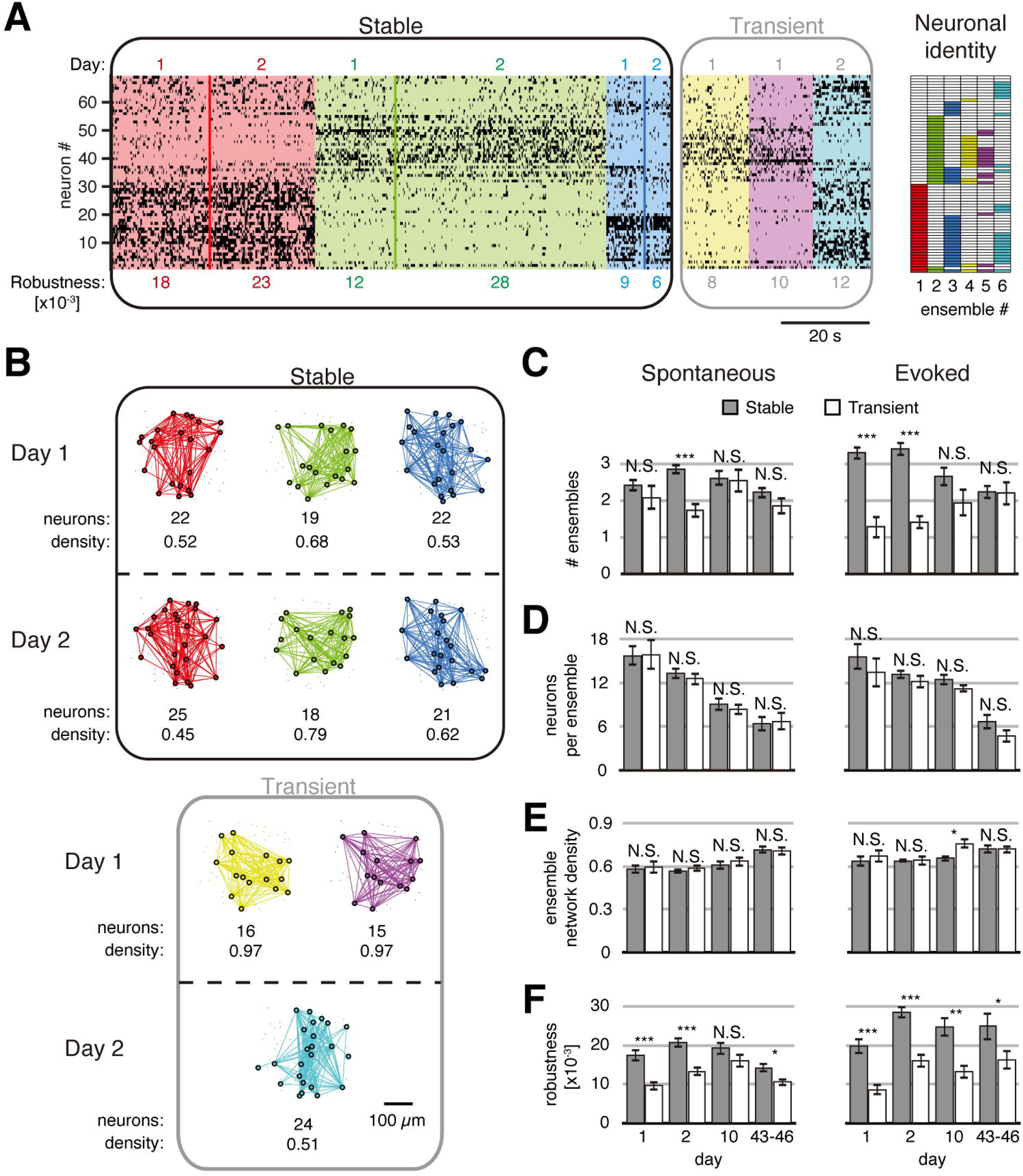
Stable and transient ensembles across days. A. Example of spontaneous ensemble activity found on a single session from day 1 and 2. Left, stable ensembles activity sorted by ensemble (background colors show different ensembles) and the day (color lines divide the activity from each day in each ensemble). Middle, transient ensembles activity sorted by day. Right, identity structure of the neuronal ensembles. Particular ensemble robustness values are at the bottom of each ensemble activity. B. Functional networks from ensembles on A preserving the spatial location of the neurons. Stable (top) and transient (bottom) ensembles separated by day observed. At the bottom of each ensemble are particular values of the number of neurons and density of the functional connectivity within the ensemble. C. Number of stable and transient ensembles during spontaneous (left) and evoked (right) activity on all days recorded. D. Count of neurons per ensemble with no significant difference between stable and transient ensembles during spontaneous (left) and evoked (right) activity on all days recorded. E. Density of the functional ensemble networks had no significant difference between stable and transient ensembles during spontaneous (left) and evoked (right) activity in almost all days, with one exception from evoked activity on day 10. F. Ensemble robustness was significantly higher in stable rather than transient ensembles during spontaneous activity (left) and evoked (right) activity in almost all days, with one exception from spontaneous activity on day 10. Mann-Whitney test: * p < 0.05, ** p < 0.01 and *** p < 0.001. See Figure 3 — Table 1.

To test if stable ensembles were a merely artifact of the analysis, we shuffled the neuronal activity of the likened session on days 1, 2, 10, 43 and 46. We found 0.8 ± 0.1 (mean ± SEM) stable ensembles from shuffled activity compared with 2.8 ± 0.1 (mean ± SEM) stable ensembles from original data (p = 1×10^−66^; Figure 3 — Figure supplement 1A). The total presence of stable ensembles during a single session of spontaneous or evoked activity was significantly higher than the ensembles found on shuffled data (75.1 ± 2.7 s and 14.4 ± 1.3 s, respectively, mean ± SEM, p = 2×10^−57^; Figure 3 — Figure supplement 1B. This suggested that our analysis of stable ensembles detected real ensembles.

We were curious to find if stable and transient neuronal ensembles had different properties. There were no significant differences between stable and transient neuronal ensembles during spontaneous or evoked activity in the number of neurons (Figure 3D) nor network ensemble density (Figure 3E). However, ensemble robustness, defined as the product of ensemble duration and similarity of its activity (see Methods), was significantly higher in stable than in transient neuronal ensembles (Figure 3F). This was the case for both spontaneous and evoked stable ensembles, indicating that stable ensembles had higher consistency in their activity.

Finally, we studied neural identity or functional connectivity of stable ensembles. We described single neuron participation on stable ensembles (Figure 4A-D). Approximately 50 % of neurons belonged to only one stable ensemble (single neurons), and less than 20 % of the neurons belonged to more than one (shared neurons), while the rest of the neurons were not part of any stable ensemble during spontaneous or evoked activity (Figure 4E). We then inquired what happened to individual neurons of stable ensembles across days. More than 60 % of neurons of a stable ensemble observed on day 1 remained in the same stable ensemble on a second session of day 1, 2, 10 and 43-46, during spontaneous and evoked activity (“maintained” neurons, Figure 4F). The rest of neurons (less than 40 %) changed their participation to another ensemble or stopped participating in detectable ensembles (“lost” neurons). Interestingly, in subsequent sessions, we found “new” neurons joining stable ensembles in a similar proportion as lost neurons. Neither maintained nor lost neurons specifically belonged to one or more than one ensemble (Figure 4G). However, functional connection density between maintained neurons from the same ensemble (0.71 ± 0.01, mean ± SEM) was significantly higher than density from lost neurons during spontaneous or evoked activity (0.35 ± 0.02, mean ± SEM, p = 7×10^−55^, Figure 4H). Therefore, poor connectivity between lost neurons could be a reason for them not being durable, and high functional connectivity indicates a possible lasting stability.

**Figure 4.**
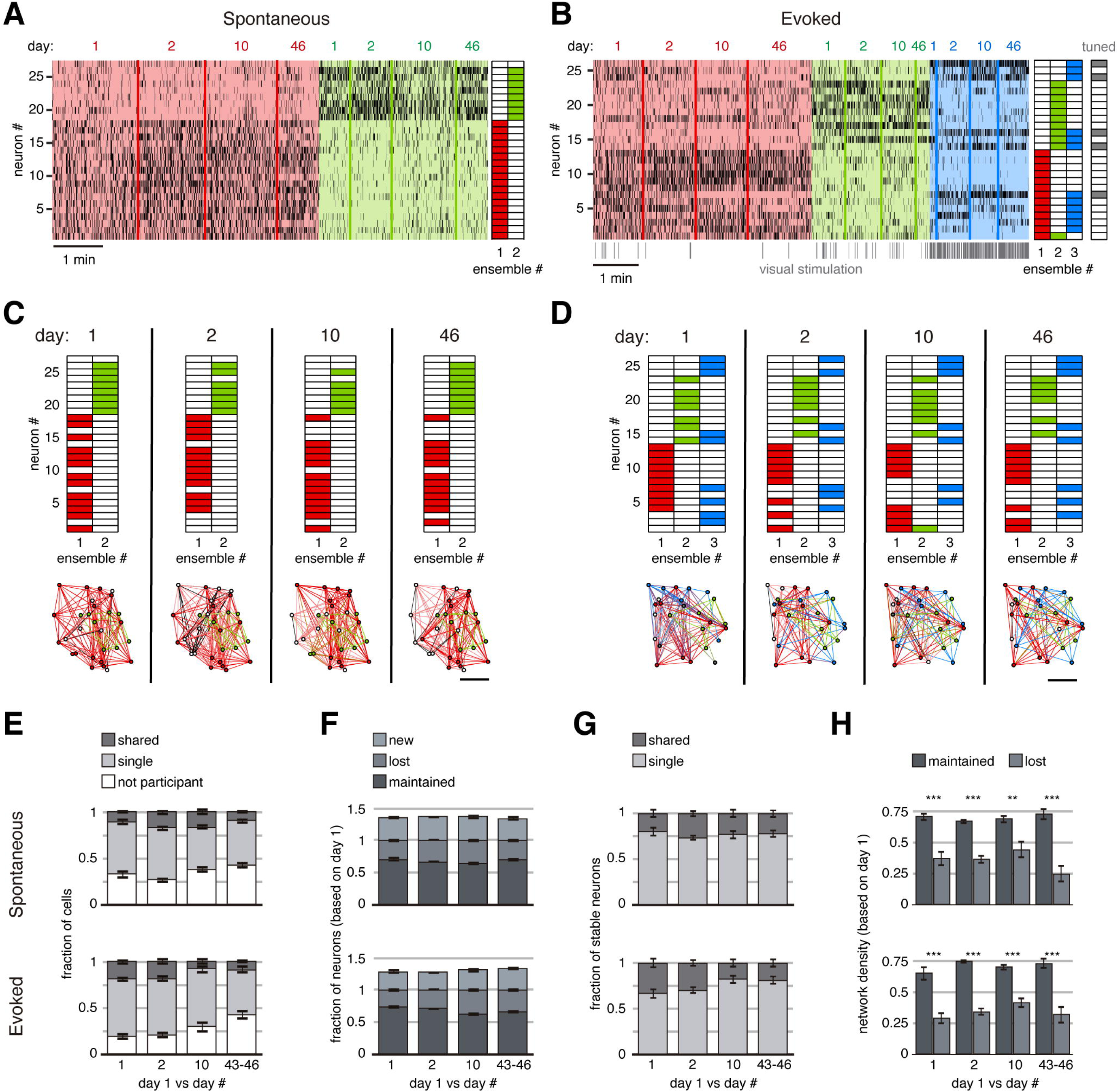
Long-term stability of spontaneous and evoked ensembles. A. Example of spontaneous ensemble activity on a single session from day 1, 2, 10 and 46. The neurons are sorted based on their ensemble identity (right). B. Example of evoked ensemble activity on a single session from day 1, 2, 10 and 46. The neurons are sorted based on their ensemble identity and it is indicated if the neurons were tuned to the visual stimulation (right). At the bottom of the raster activity are marked the first ∼500 ms (of the 2 s) of every visual stimulation (50 per session). Note there is a particular stable neuronal ensemble (blue ensemble) mainly evoked at the onset of the visual stimulation. C. Neuronal ensembles separated by days (identified independently). Top, the neuronal identity sorted as in A. Bottom, functional network plotted preserving their neuronal spatial locations. Colors indicate the ensemble which neurons belonged, white color indicates no participation in any stable ensemble. Scale bar: 50 µm. D. Neuronal ensembles separated by days (identified independently). Top, the neuronal identity sorted as in B. Bottom, functional network plotted preserving their neuronal spatial locations. Colors indicate the ensemble which neurons belonged, white color indicates no participation in any stable ensemble, gray color indicates participation in more than one stable ensemble. Scale bar: 50 µm. E. Fraction of neurons during spontaneous (top) or evoked (bottom) activity from day 1, 2, 10 and 43-46 which participated in two or more stable ensembles (shared), only one (single) and without participation in a stable ensemble. F. Fraction of neurons during spontaneous (top) and evoked (bottom) activity from day 1, 2, 10 and 43-46 which remained in the same ensemble (maintained), changed their ensemble or stopped to participate (lost) and new neurons. G. Fraction of stable neurons during spontaneous (top) and evoked (bottom) activity from day 1, 2, 10 and 43-46 which participated in one stable ensemble (single) o more (shared). H. Network density within stable ensembles during spontaneous (top) and evoked (bottom) activity was significantly higher in maintained neurons than lost neurons on all days 1, 2, 10 and 43-46 (p < 0.01). Density was computed from functional connectivity analyzed on day 1. Data are presented as mean ± SEM. Mann-Whitney test: ** p < 0.01 and *** p < 0.001. See Figure 4 — Table 1.

## Discussion

Starting with Wiesel and Hubel’s landmark studies in the 1960’s, the plasticity and stability of neuronal activity of the visual cortex has been explored in single cell studies, characterizing single neuron properties such as receptive fields, tuning, spiking rate, etc. (Wiesel and Hubel, 1963; Wandell and Smirnakis, 2009; Ranson, 2017; Jeon et al., 2018). Our results, using multineuronal imaging methods, and a microcircuit perspective (Shepherd & Grillner, 2010; Yuste, 2015; Bargas and Pérez-Ortega, 2017), complement this field by measuring emergent properties such as functional connectivity, neuronal ensembles, network properties, robustness and stability. In spite of the incomplete sampling of circuit activity and potential alterations on the circuit due to repeated experiments, our results indicate that some neuronal ensembles are robust and can last several weeks. From all the neuronal ensemble properties we measured, there was no relevant difference between spontaneous and evoked activity, consistent with the hypothesis that they are the same and that sensory stimulus reactivate existing ensembles.

Volumetric two-photon calcium imaging allowed carefully tracking the same neurons and identifying neuronal ensembles across days. Nevertheless, our experiments had limitations, as imaging data of animals became noisier across time goes (Figure 1G), likely due to accumulated experimental issues, which made us difficult to track every neuron with sufficient signal to noise (PSNR > 18 dB, Figure 1H). Nevertheless, neurons tracked for weeks remained at the same level of activity as on the first day (Figure 1I). Interestingly, even after 5 minutes of recording, only ∼50 % of previously active neurons continued to be active. This, and the fact that lost neurons in ensembles are replaced by new ones (Figure 4H), is consistent with the possibility that the circuit maintains a basic homeostatic level of activity, with neurons rotating in and out, while the ensembles are maintained. Thus, one could posit that one of the functions of neuronal ensembles is precisely to maintain a stable functional state in the midst of an ongoing homeostatic replacement of the activity of individual neuronal elements. At the same time, some neurons appeared particular stable, and at least 60 % of the neurons belonging to the same ensemble remain on the same ensemble 5 min and up to 46 days later (Figure 4F). Since maintained neurons in stable ensembles appear to have stronger functional connectivity, it is possible that they represent a special cell type, or a class of neuron with special functional properties, forming the core of an ensemble.

In summary, we found a lasting permanence of neuronal ensembles in 2/3 layer of mouse visual cortex. The coactivation of a group of neurons is at least partly explained by its stronger internal functional connectivity, mediated by short- or long-term synaptic plasticity (Carrillo-Reid et al., 2015b; Hoshiba et al., 2017), which is thought basis of learning and memory. Because of this, we hypothesize that stable neuronal ensembles implement long-term memories whereas transient ensembles could represent the emergence of new memories or the degradation of existing ones.

## Methods

### Animals

Experiments were performed on transgenic mice Vglut1-Cre crossed with TIGRE2.0 lines Ai162 (TIT2L-GC6s-ICL-tTA2) or Ai148 (TIT2L-GC6f-ICL-tTA2). Mice were housed on a 12 h light-dark cycle with food and water *ad libitum*. Head-plate procedure and cranial window was executed after 50 days of age. Mice health was checked daily. All experimental procedures were carried out in accordance with the US National Institutes of Health and Columbia University Institutional Animal Care and Use Committee.

### Head-plate procedure and cranial window

Adult transgenic mice GCaMP6s (n = 4) and GCaMP6f (n = 2) were anesthetized with isoflurane (1.5 - 2 %). Body temperature was maintained at 37 °C with a heating pad and eyes were moisturized with eye ointment. Dexamethasone sodium phosphate (0.6 mg/kg) and enrofloxacin (5 mg/kg) were administered subcutaneously. Carprofen (5 mg/kg) was administered intraperitoneally. A custom designed titanium head-plate was attached to the skull using dental cement. Then, a craniotomy was made of 3 mm in diameter with center at 2.1 mm lateral and 3.4 mm posterior from bregma. A 3-mm circular coverslip was implanted and sealed using cyanoacrylate and cement. After surgery animals received carprofen injections for 2 days as post-operative pain medication. Mice were allowed to recover for 5 days with food and water *ad libitum*.

### Visual stimulation

Visual stimuli were generated using a custom-made app on MATLAB (https://www.mathworks.com/matlabcentral/fileexchange/78670-drifting-gratings-generator-for-visual-stimulation) displaying on an LCD monitor positioned 15 cm from the right eye at 45 degrees to the long axis of the animal. The red and green channels of the monitor were disabled to avoid light contamination in the imaging photomultiplier (PMT), only blue channel was enabled. We used two protocols to display in the monitor. The first was in absence of visual stimulation, the monitor was displaying a static blue screen, we used it to record spontaneous activity during 5 min per session. The second protocol was for visual stimulation consisting of a full sinusoidal gratings (100 % contrast, 0.13 cycles/deg, 5 cycles/s) drifting in a single direction per mouse (0° or 270°) presented for 2 s, followed by a random amount between 1 to 5 s of mean luminescence. The visual stimulus is presented 50 times during 5 min per session. We performed 3 consecutive sessions (5 min apart) per protocol per day of experiment. See Figure 1 — Table 1 for a detailed sessions recorded per mouse.

### Volumetric two-photon calcium imaging

Imaging experiments were performed from 20-150 days after head-plate procedure. Each mouse was placed on a treadmill with its head fixed under the two-photon microscope (Ultima IV, Bruker). Animals were acclimated to the head restraint for periods between 5 to 15 min for at least 2 days, and exposed to visual stimulation sessions before the recordings presented here. The imaging setup was completely enclosed with blackout fabric to avoid light contamination leaking into the PMT. An imaging laser (Ti:sapphire, λ = 920 nm, Chameleon Ultra II, Coherent) was used to excite a genetically encoded calcium indicator (GCaMP6s or GCaMP6f). The laser beam on the sample (30-60 mW) was controlled by a high-speed resonant galvanometer scanning an XY plane (256×256 pixels) at 17.7 ms (frame period) covering a field of view of 312×312 µm using a 25X objective (NA 1.05, XLPlan N, Olympus). An electrically tunable lens (ETL) was used to change the focus (z-axis) during the recording. We recorded consecutively 3 planes at different depths (−5, 0 and 5 µm from a reference z-axis) waiting 9.3 ms between planes for ETL to stabilize the focus. Thus, we collected three frames, one per depth, every 81 ms for 5 min (single session, 3,704 frames per plane). Imaging was controlled by Prairie View and ETL was synchronized using a DAQ (USB-6008, NI) controlled by a custom-made app on MATLAB (https://www.mathworks.com/matlabcentral/fileexchange/78245-etl-controller-for-volumetric-imaging).

### Recording same neuronal region through days

In the first day of experiment, we recorded the vascularization of the pia at 10X and 25X using bright-light microscopy. We fixed the depth to 140 µm from pia to record a reference image (calcium imaging) and a second reference image of the center of the field of view using an extra 4X optical zoom (day 1 in Figure 1G). We carefully preserved unchanged the position of the microscope and the base we placed the mice. For following days of recording, we looked for matching the reference image of vascularization at 10X, then 25X. After that, we looked 140 µm depth from pia trying to match the reference image on *x* and *y* axis, then we used a 4X extra optical zoom to finely match the second reference image on z axis (day 2, 10 and 43 or 46 in Figure 1G). We performed a volumetric imaging to record 3 planes: one reference plane, one 5 µm above and one 5 µm below in order to emend tilt, which in our system does not go beyond 5 µm. We extracted the maximum intensity projection from the 3 planes resulting in a single video for each session (Figure 1D).

### Neuronal activity extraction

We used a custom-made graphical user interface (GUI) on MATLAB (https://github.com/PerezOrtegaJ/Catrex_GUI) to extract the binary raster activity from every single session video (5 min, 3,704 frames). First, we performed a non-rigid motion correction taking as a reference the mean of the 185 frames (5 %) with less motion artifacts. Then, we searched the ROIs with a modified version of Suite2P algorithm (Pachitariu et al., 2016, Figure 1E). ROIs were preserved if they fulfill the following criteria (fixing radius to 4 µm): 0.5*π *radius^2 < area < 4*π*radius^2; roundness > 0.2; perimeter < 3.5*π*radius; eccentricity < 0.9 and overlapping < 60 %. Calcium signal from each ROI was extracted measuring the changes in fluorescence with respect of its local neuropil (F_raw_ — F_n_) / F_n_, where F_raw_ is the signal from the ROI and F_n_ is the signal of its local neuropil 10 times ROI radius. ROI local neuropil is not including signal from ROIs if presented within the area. Then we computed the peak signal-to-noise ratio PSNR = 20 · log10(max(F_raw_ — F_n_) / σ_n_), where max represents a maximum function, and σ_n_ represents the standard deviation of the local neuropil. We evaluated the ROIs again keeping them if PSNR > 18 dB. Then we smoothed the calcium signal with a 1 s window average to perform a spike inference using the foopsi algorithm (Friedrich & Paninski, 2016). We binarized the spike inference signal, placing 1 if there was spikes inferred and 0 if not. We placed all binarized signals from every ROI in a N x F raster matrix, where N is the number of active neurons and F the number of frames. This matrix is visualized as a raster plot, where ones in the matrix are the dots representing the active frames of the neurons (Figure 1F).

### Tracking neurons across days

We computed a rigid and then a non-rigid motion correction between the binary image of the ROIs shape between a single session of day 1 and a single session from day 1, 2, 10 and 43 or 46 (Figure 1J). Then, we looked for the intersection (in pixels) between ROIs of the neurons from two sessions (intersection > 0.5* π *radius^2) and evaluate the Euclidean distance between centroids of the ROIs intersected keeping it if the distance < radius. We used the raster matrix only with the tracked neurons between sessions.

### Identification of neuronal ensembles based on functional connectivity

To analyze neuronal ensembles from raster activity we used a custom-made GUI on MATLAB (https://github.com/PerezOrtegaJ/Neural_Ensemble_Analysis). Functional connectivity represents the significant coactivity between every pair of neurons from a raster matrix. The number of coactivations Co_ab_ between neuron *a* and *b* was computed counting in how many single frames were both neurons simultaneously active. To identify significance, we generated 1,000 surrogates of neurons *a* and *b* by random circular shifting their activity in time to disrupt their temporal dependency. We counted the number of surrogate coactivations S_ab,i_ in each iteration *i*, building a cumulative distribution of S_ab_ selecting a threshold T_ab_ of coactivations at the 95%. If the actual number of coactivations Co_ab_ is above threshold T_ab_ we put a functional connection between neuron *a* and *b* (Figure 2B). Doing this with every pair of neurons we got a functional neuronal network, where every node is a neuron and every link represents a significant coactivity between them (Figure 2C). We used the functional connectivity to rebuild the raster matrix to keep the significant coactivity of the neurons. To do so, we identified the active neurons of every single frame, we looked to their functional connectivity, if a neuron has no connection its activity was removed from that frame. At the end, we also removed the frames with less than 3 coactive neurons (Figure 2D). Then we computed the Jaccard similarity between all single frames (column vectors) of the rebuilt raster matrix. A hierarchical clustering tree with single linkage was obtained to identify the more similar vectors by keeping the branch with more than 2/3 of Jaccard similarity (Figure 2E). With the more similar vectors we performed a hierarchical clustering with Ward linkage and grouped based on a contrast index (Beggs & Plenz, 2004). Each group of column vectors is the activity of the neuronal ensemble (Figure 2F), i.e. the raster matrix E_j_ of an ensemble *j* of size N x F_j_, where N is the number of neurons and F_j_ is number of frames where the ensemble *j* was active.

### Demixing neuronal ensemble identity

The neuronal ensemble activity E_j_ was used to identify the participation of each neuron in the ensemble *j*. We computed the functional connectivity similarly as described above, but incorporating the correlation of the neurons between the times where the ensemble was active. To do so, we got a binary vector V_j_ representing the times were the ensemble *j* was active (1) or not (0). Vector V_j_ was of size 1 x F, where F is the number of frames of the session. A Pearson correlation coefficient P_j,a_ between vector V_j_ and the activity of neuron *a* was computed. Then we got an ensemble weight W_j,ab_ between neurons *a* and *b* in ensemble *j*, which integrates their correlation with the ensemble *j* and their number of coactivations as follows W_j,ab_ = P_j,a_ · P_j,b_ · Co_ab_. To identify significance, we generated 1,000 surrogates of neurons *a* and *b* shuffling their activity as described before, and assigning randomly a value from the correlation with the ensemble *j* (P_j_). Then we compute the surrogate weight SW_j,ab,i_ in each iteration *i*, building a cumulative distribution of SW_j,ab_ selecting a threshold TW_j,ab_ of coactivations at the 95%. If the actual ensemble weight SW_j,ab_ is above threshold TW_j,ab_ we put a functional connection between neuron *a* and *b*. A neuron is considered to be part of an ensemble if at least had one single functional connection (Figure 2G). A neuron could be part of more than one neuronal ensemble (shared), only one ensemble (single) or in any ensemble (not participant).

### Comparing neuronal ensembles between following sessions

Taking the first session on day 1 and a second session from day 1, 2, 10, 43 or 43, we got the raster matrix from each session with only tracked neurons (same active neurons in both sessions). Neuronal ensembles were extracted from each raster matrix independently. If there were 50 % or more neurons in an ensemble *j* from the first session belonging to an ensemble *k* from the second session we called it “stable” ensemble, otherwise we called it “transient” ensemble.

### Ensemble measures

*Ensemble network density*, fraction of present functional connections to possible connections within an ensemble. *Ensemble robustness*, we introduced here as *robustness* = *similarity* · *activity*, where similarity is the average of the Jaccard similarity between every pair of column vectors of the ensemble matrix raster E_j_, and activity is the fraction of ensemble active frames to the total frames of the session. The higher the value, the higher the robustness. *Stability of neurons*: given a stable ensemble, a “maintained” neuron participated during the first session on day 1 and during a second session on the following days. A “lost” neuron participated only in the first session but not in a second session, and a “new” neuron did not participate in the first session but participate in a second session. The fraction is based on total neurons in an ensemble from day 1. *Promiscuity of neurons*: “shared” neurons is the fraction of neurons participating in more than one ensemble; “single” neurons participate in only one ensemble; and “not participant” neurons do not belong to any ensemble. *Tuned neurons*: we consider a tuned neuron if its number of active frames during visual stimulation was significantly higher than its number of active frames during periods with no visual stimulation (P < 0.5, t-test).

## Acknowledgments

We thank James Holland for his assistance and members of the Yuste Lab for useful comments. This project was supported by R01EY011787 and R01MH115900. R.Y. is an Ikerbasque Research Professor at the Donostia International Physics Center (DIPC). J.P. has a postdoctoral fellowship from the National Council of Science and Technology from Mexico (CONACYT). The authors have no competing financial interests to declare. J.P., T.A. and R.Y. conceived the project, planned experiments and discussed results. T.A. and J.P. performed surgeries, J.P. performed experiments, coded the software and analyzed the data. J.P. and R.Y. wrote the paper. R.Y. assembled and directed the team and secured funding and resources. J.P. dedicates this paper to the memory of Amparo Rodríguez-Cruz.

## Figure Legends

**Figure 1 — Figure supplement 1. Active neurons in different session times**.

A. Total number of active neurons (mean ± SEM, n = 2 mice) between 2 sessions on the same day with equal duration, sessions started from 10 s to 10 min. An exponential function was fitted (R^2^ = 0.99). Note that number of active neurons started a plateau around 5 min of recording length.

B. Percentage of common active neurons between the sessions showed on A. An exponential function was fitted (R^2^ = 0.77) and started a plateau around 3 min increasing slowly as sessions length increase.

**Figure 1 — Table 1. Mice and recording days**.

**Figure 1 — Table 2. Neuronal activity across days**.

**Figure 2 — Figure supplement 1. Number of ensembles in spontaneous and evoked activity**.

A. Distribution of the number of ensembles found during spontaneous (top) and evoked activity (bottom) across days. No significant differences were found between day 1 and the following days during spontaneous and evoked activity (day 2, p = 0.97 and p = 0.86; day 10, p = 0.41 and p = 0.99; and day 43-46 (p = 0.60 and p = 0.99, respectively; Kruskal-Wallis test, post hoc Tukey-Kramer).

**Figure 3 — Table 1. Statistics of stable and transient ensembles**.

**Figure 3 — Figure supplement 1. Shuffling controls**.

A. Distribution of the number of ensembles found during the real activity versus shuffled activity.

B. Distribution of the length of time the ensembles found were present during the actual activity versus the expected by chance shuffling the activity.

C. Number of ensembles found during spontaneous (top) and evoked (bottom) activity through the days compared with the shuffled version (P < 0.001 for all comparisons; Mann-Whitney test).

D. Length of time the ensembles were present during spontaneous (top) and evoked (bottom) activity through the days compared with the shuffled version (P < 0.001 for all comparisons; Mann-Whitney test).

**Figure 3 — Video supplement 1. Stable ensembles in spontaneous activity on day 1 and 46**.

Raw videos of 2 min length (accelerated 5x) from day 1 (top-left) and day 46 (top-right). Circles are identifying neurons from stable neuronal ensembles, colors represent different ensembles. Scale bar: 50 µm. At the bottom, summation of calcium signals from each ensemble.

**Figure 3 — Video supplement 2. Stable ensembles in evoked activity on day 1 and 46**.

Raw videos of 2 min length (accelerated 5x) from day 1 (top-left) and day 46 (top-right). Circles are identifying neurons from stable neuronal ensembles, colors represent different ensembles. Scale bar: 50 µm. At the bottom, summation of calcium signals from each ensemble.

**Figure 4 — Table 1. Neuronal composition of stable ensembles**.

## Notes

https://www.mathworks.com/matlabcentral/fileexchange/78670-drifting-gratings-generator-for-visual-stimulation

https://www.mathworks.com/matlabcentral/fileexchange/78245-etl-controller-for-volumetric-imaging

https://github.com/PerezOrtegaJ/Catrex_GUI

https://github.com/PerezOrtegaJ/Neural_Ensemble_Analysis

### Competing Interest Statement

The authors have declared no competing interest.

