## Supplementary figures and images for "Long-term stability of neuronal ensembles in mouse visual cortex"

### Figure 1 - Figure supplement 1

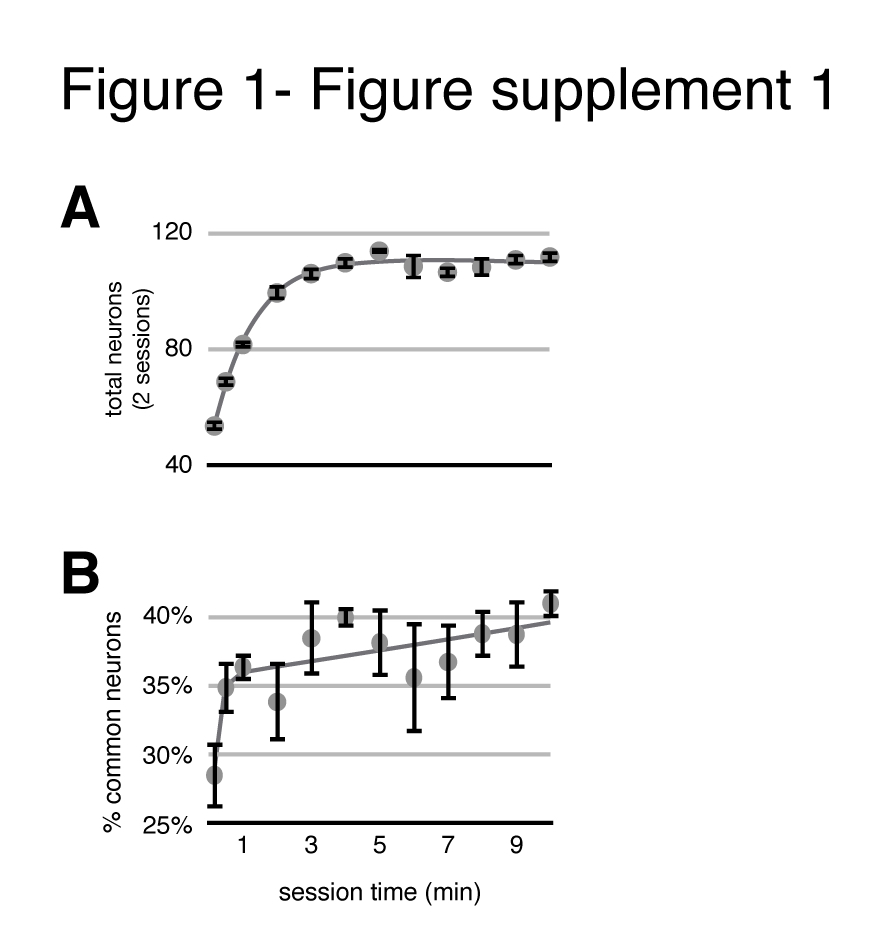

### Figure 2 - Figure supplement 1

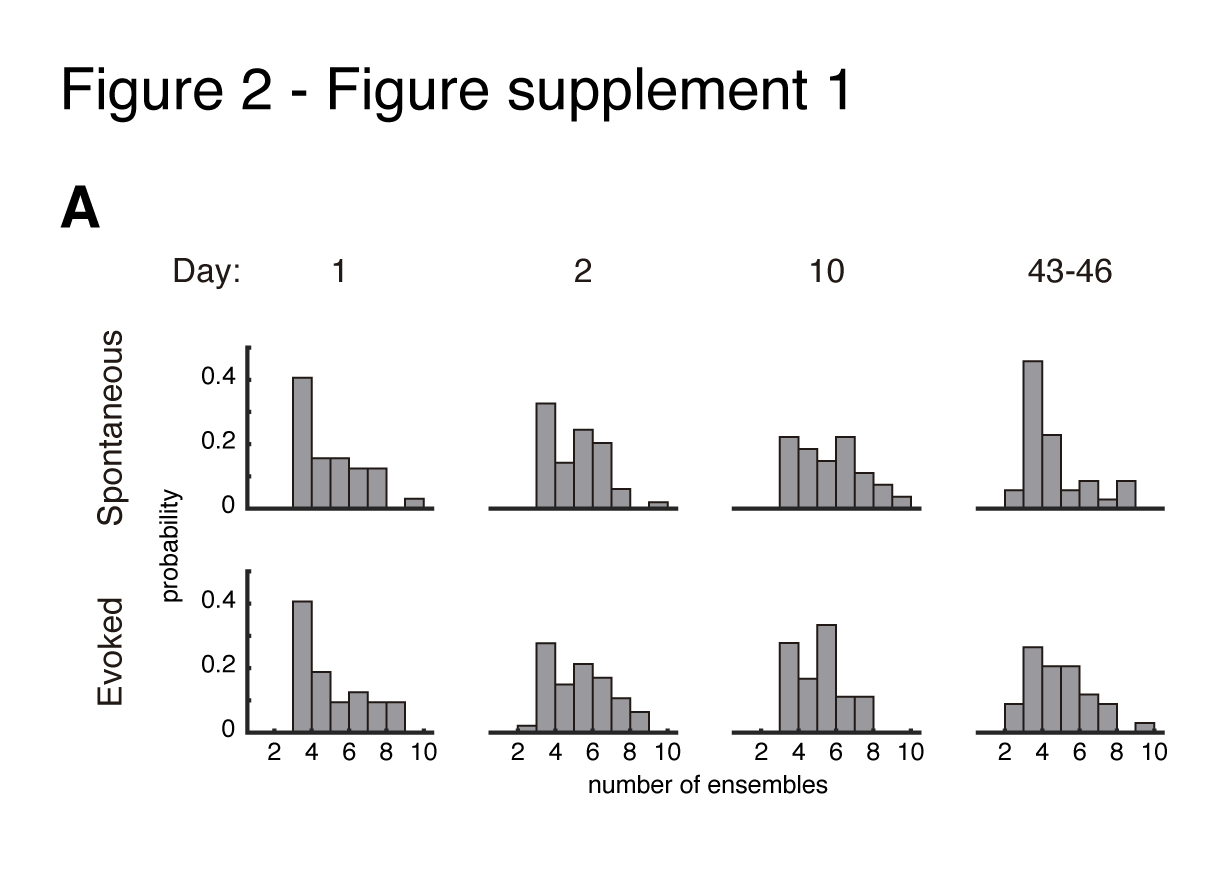

### Figure 3 - Figure supplement 1

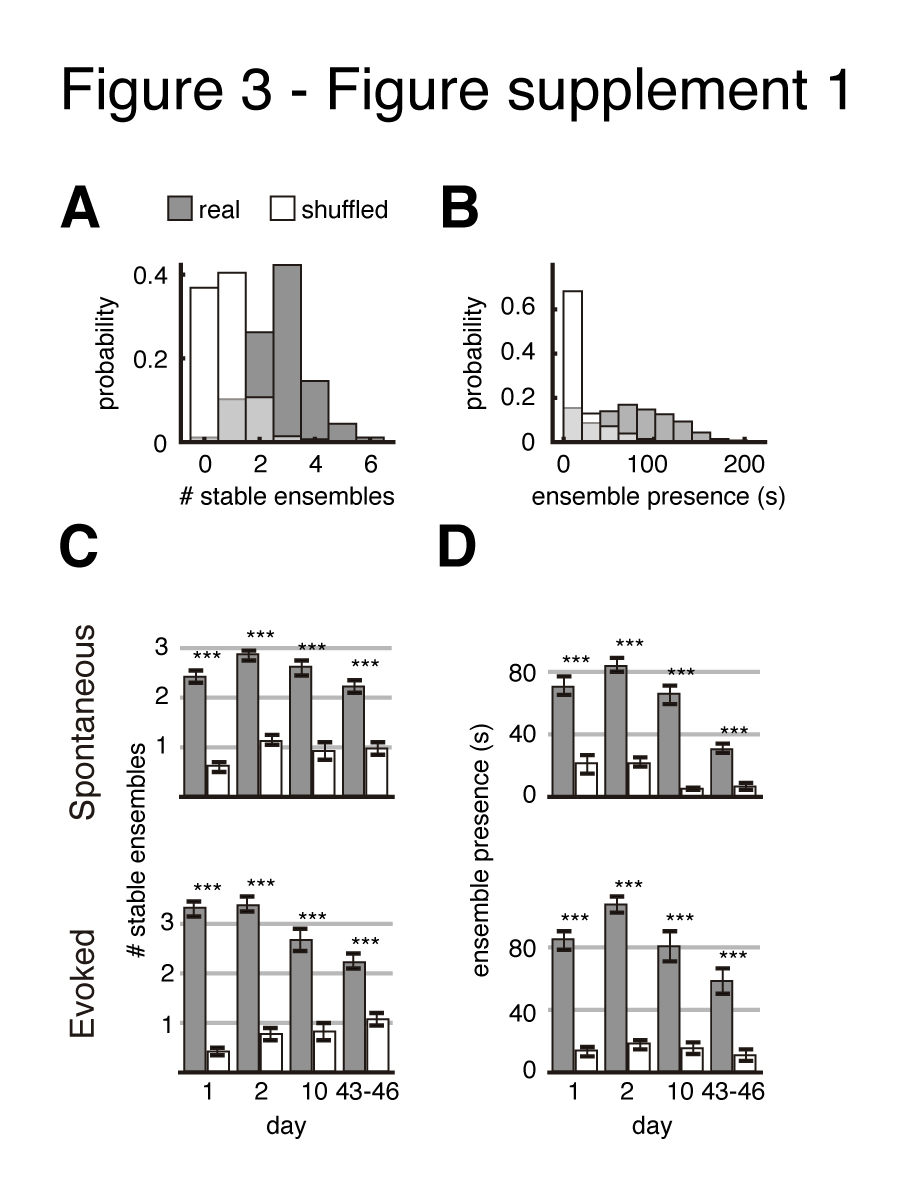
