## Supplementary material for "Long-term stability of neuronal ensembles in mouse visual cortex": Figure 1 - Table 1

**Figure 1 — Table 1. Mice and recording days.**

| **Mouse #** | **Sex** | **GCaMP** | **PD at day 1** | **Spontaneous (day)** | | | | | **Visual stimulation (day)** | | | | |
| --- | --- | --- | --- | --- | --- | --- | --- | --- | --- | --- | --- | --- | --- |
|  |  |  |  | **1** | **2** | **10** | **43** | **46** | **1** | **2** | **10** | **43** | **46** |
| **1** | ♂️ | 6s | 98 | x | x |  | x |  | x | x |  | x |  |
| **2** | ♂️ | 6s | 90 | x* | x* |  |  |  | x* | x* |  |  |  |
| **3** | ♂️ | 6s | 90 | x | x |  | x |  | x | x |  | x |  |
| **4** | ♀️ | 6s | 197 | x | x | x |  |  | x | x |  |  |  |
| **5** | ♂️ | 6f | 133 | x | x | x |  | x | x | x | x |  | x |
| **6** | ♂️ | 6f | 133 | x | x | x |  | x | x | x | x |  | x |
| n  = | | | | 17 | 17 | 9 | 12 | | 17 | 17 | 6 | 12 | |
| x = 3 sessions; x* = 2 sessions | | | |  |  |  |  | |  |  |  |  | |
