## Supplementary material for "Long-term stability of neuronal ensembles in mouse visual cortex": Figure 1 - Table 2

**Figure 1 — Table 2. Neuronal activity across days.**

|  |  | **day 1** | **day 2** | **p**  **days 1 VS 2** | **day 10** | **p**  **days 1 VS 10** | **day 43-46** | **p**  **days 1 VS 43-46** |
| --- | --- | --- | --- | --- | --- | --- | --- | --- |
| **Active neurons number** | spontaneous | 83 ± 6 | 78 ±  6 | 1 | 70 ± 9 | 0.64 | 51 ± 10 | ***0.023** |
|  | evoked | 85 ± 5 | 80 ±  6 | 0.84 | 87 ± 9 | 1 | 51 ± 10 | ***0.013** |
| **% time single neuron is active** | spontaneous | 16 ± 1.1 % | 15 ± 0.9 % | 0.60 | 17 ± 0.4 % | 0.15 | 15 ± 1.3 % | 0.76 |
|  | evoked | 14 ± 0.9 % | 15 ± 0.5 % | 1 | 16 ± 0.4 % | ***0.02** | 15 ± 0.7 % | 0.90 |
| **% common neurons**  **(day 1 VS day #)** | spontaneous | 42 ± 1.9 % | 33 ± 0.9 % | 0.10 | 22 ± 1.2 % | *****2x10^-6^** | 15 ± 1.5 % | *****4x10^-9^** |
|  | evoked | 40 ± 3.7 % | 34 ± 1.3 % | 0.26 | 25 ± 0.5 % | *****2x10^-4^** | 15 ± 1.5 % | *****4x10^-9^** |
| Data are presented as mean ± SEM. Kruskal-Wallis test with post hoc Tukey-Kramer: * p < 0.05, ** p < 0.01 and *** p < 0.001 | | | | | | | | |
