## Supplementary material for "Long-term stability of neuronal ensembles in mouse visual cortex": Figure 3 - Table 1

**Figure 3 — Table 1. Statistics of stable and transient ensembles.**

|  |  | **Number of ensembles** | | | **Neurons per ensemble** | | | **Density** | | | **Ensemble robustness** | | |
| --- | --- | --- | --- | --- | --- | --- | --- | --- | --- | --- | --- | --- | --- |
|  | **day** | **stable** | **transient** | **p** | **stable** | **transient** | **p** | **stable** | **transient** | **p** | **stable** | **transient** | **p** |
| **Spontaneous** | 1 | 2.4 ± 0.2 | 2.1 ± 0.3 | 0.33 | 16 ± 1.4 | 16 ± 2.0 | 0.61 | 0.58 ± 0.03 | 0.60 ± 0.04 | 0.54 | 17 ± 1.5 | 10 ± 1.0 | *****1x10^-5^** |
|  | 2 | 2.9 ± 0.1 | 1.7 ± 0.2 | *****1x10^-5^** | 13 ± 0.7 | 13 ± 0.8 | 0.30 | 0.57 ± 0.01 | 0.59 ± 0.02 | 0.99 | 21 ± 1.1 | 13 ± 1.0 | *****5x10^-5^** |
|  | 10 | 2.6 ± 0.2 | 2.6 ± 0.3 | 0.50 | 9 ± 0.9 | 8 ± 0.7 | 0.47 | 0.61 ± 0.03 | 0.63 ± 0.03 | 0.92 | 19 ± 1.5 | 16 ± 1.7 | 0.25 |
|  | 43-46 | 2.2 ± 0.1 | 1.9 ± 0.2 | 0.10 | 6 ± 1.0 | 7 ± 1.2 | 0.93 | 0.72 ± 0.03 | 0.71 ± 0.03 | 0.89 | 14 ± 1.0 | 11 ± 0.8 | ***0.04** |
| **Evoked** | 1 | 3.3 ± 0.2 | 1.3 ± 0.3 | *****5x10^-5^** | 16 ± 1.7 | 13 ± 2.0 | 0.45 | 0.64 ± 0.03 | 0.68 ± 0.04 | 0.53 | 20 ± 1.9 | 9 ± 1.3 | *****6x10^-6^** |
|  | 2 | 3.4 ± 0.2 | 1.4 ± 0.2 | *****7x10^-11^** | 13 ± 0.6 | 12 ± 0.8 | 0.35 | 0.64 ± 0.01 | 0.64 ± 0.03 | 0.79 | 29 ± 1.5 | 16 ± 1.5 | *****2x10^-7^** |
|  | 10 | 2.7 ± 0.2 | 1.9 ± 0.4 | 0.13 | 12 ± 0.8 | 11 ± 0.5 | 0.26 | 0.66 ± 0.02 | 0.76 ± 0.03 | ***0.02** | 25 ± 2.5 | 13 ± 1.7 | ****3x10^-3^** |
|  | 43-46 | 2.2 ± 0.2 | 2.2 ± 0.3 | 0.46 | 7 ± 0.9 | 5 ± 0.8 | 0.19 | 0.72 ± 0.2 | 0.72 ± 0.03 | 0.80 | 25 ± 3.4 | 16 ± 2.3 | ***0.01** |
| Data are presented as mean ± SEM. Mann-Whitney test: * p < 0.05, ** p < 0.01 and *** p < 0.001. | | | | | | | | | | | | | |
