## Supplementary material for "Long-term stability of neuronal ensembles in mouse visual cortex": Figure 4 - Table 1

**Figure 4 — Table 1. Neuronal composition of stable ensembles.**

|  |  | **Fraction of neurons** | | | **Fraction of neurons** | | | **Fraction of stable neurons** | | **Network density on day 1** | | |
| --- | --- | --- | --- | --- | --- | --- | --- | --- | --- | --- | --- | --- |
|  | **day 1**  **vs**  **day #** | **single** | **shared** | **non participant** | **maintained** | **lost** | **new** | **single** | **shared** | **maintained** | **lost** | **p** |
| **Spontaneous** | 1 | 0.57 ± 0.03 | 0.1 ± 0.02 | 0.33 ± 0.04 | 0.7 ± 0.02 | 0.3 ± 0.02 | 0.35 ± 0.02 | 0.8 ± 0.04 | 0.2 ± 0.04 | 0.71 ± 0.03 | 0.38 ± 0.05 | *****4x10^-5^** |
|  | 2 | 0.56 ± 0.02 | 0.17 ± 0.02 | 0.26 ± 0.02 | 0.67 ± 0.01 | 0.33 ± 0.01 | 0.36 ± 0.02 | 0.74 ± 0.03 | 0.26 ± 0.03 | 0.67 ± 0.02 | 0.37 ± 0.03 | *****1x10^-10^** |
|  | 10 | 0.46 ± 0.03 | 0.16 ± 0.03 | 0.38 ± 0.03 | 0.63 ± 0.01 | 0.37 ± 0.02 | 0.37 ± 0.02 | 0.77 ± 0.04 | 0.23 ± 0.04 | 0.69 ± 0.03 | 0.44 ± 0.07 | ****0.003** |
|  | 43-46 | 0.48 ± 0.02 | 0.1 ± 0.02 | 0.42 ± 0.03 | 0.7 ± 0.02 | 0.3 ± 0.02 | 0.34 ± 0.02 | 0.78 ± 0.04 | 0.22 ± 0.04 | 0.73 ± 0.04 | 0.25 ± 0.07 | *****3x10^-6^** |
| **Evoked** | 1 | 0.62 ± 0.02 | 0.19 ± 0.02 | 0.19 ± 0.02 | 0.73 ± 0.02 | 0.27 ± 0.02 | 0.28 ± 0.03 | 0.67 ± 0.05 | 0.33 ± 0.05 | 0.65 ± 0.05 | 0.29 ± 0.04 | *****2x10^-4^** |
|  | 2 | 0.61 ± 0.03 | 0.19 ± 0.03 | 0.2 ± 0.03 | 0.71 ± 0.01 | 0.29 ± 0.01 | 0.27 ± 0.01 | 0.7 ± 0.04 | 0.30 ± 0.04 | 0.75 ± 0.01 | 0.34 ± 0.03 | *****3x10^-15^** |
|  | 10 | 0.63 ± 0.04 | 0.08 ± 0.03 | 0.29 ± 0.05 | 0.62 ± 0.01 | 0.38 ± 0.01 | 0.31 ± 0.02 | 0.82 ± 0.04 | 0.18 ± 0.04 | 0.7 ± 0.02 | 0.42 ± 0.04 | *****6x10^-6^** |
|  | 43-46 | 0.49 ± 0.04 | 0.09 ± 0.02 | 0.42 ± 0.04 | 0.67 ± 0.02 | 0.33 ± 0.02 | 0.34 ± 0.02 | 0.81 ± 0.04 | 0.19 ± 0.04 | 0.73 ± 0.04 | 0.32 ± 0.06 | *****9x10^-6^** |
| Data are presented as mean ± SEM. Mann-Whitney test: * p < 0.05, ** p < 0.01 and *** p < 0.001. | | | | | | | | | | | | |
